# Lysosome-like structures in the nucleolus mediate H2Aub degradation under glucose starvation

**DOI:** 10.1101/2025.06.30.662255

**Authors:** Jinli Jin, Senyu Ma, Jing Shi, Hanghang Lou, Yunxiao Li, Yao Yao, Na Fang, Yuqi Liu, Lu Yang

**Affiliations:** Department of Pharmacology, School of Basic Medical Sciences, Henan University, Kaifeng, Henan, 475004, China; Naval Medical Center, Naval Medical University, Shanghai, 200433, China; National Key Laboratory of Immunity and Inflammation, Shanghai, 200433, China; Translational Medical Research Center, Naval Medical University, Shanghai, 200433, China; Shanghai Baimu Biotechnology Co., Ltd., Room 1205, No. 968 Yierba Memorial Road, Baoshan District, Shanghai, 200435, China; Cardiac Department, Sixth Center of Chinese PLA General Hospital; Cardiac Department, First Center of Chinese PLA General Hospital; National Key Laboratory of Kidney Diseases; Beijing 100141, China

## Abstract

Glucose starvation leads to the degradation of intracellular proteins to generate amino acids for energy production. However, it remains unclear which proteins undergo rapid degradation to supply amino acids as energy sources during glucose starvation. To investigate the molecular mechanism underlying the transition of carbon source from glucose to proteins under glucose starvation, we first employed quantitative proteomics to examine the impact of glucose deprivation on protein levels. We discovered that histone degradation produces the highest amount of glutamine during glucose starvation. Further investigation revealed that histone H2A undergo degradation in the nucleolus in the form of mono-ubiquitination (H2Aub). The nucleolus contains lysosome-like structures that degrade monoubiquitinated histones, potentially participating in the carbon source shift from glucose to amino acids for energy production. Through preliminary screening, we identified Midnolin as the receptor for H2Aub, mediating its degradation in nucleolar lysosome-like structures. Dysregulation of carbon source transition from glucose to proteins, through knockdown of Midnolin results in cell cycle arrest, affecting cellular growth and proliferation. In conclusion, our findings demonstrate that nucleolar lysosome-like structures degrading monoubiquitinated histones exist during glucose starvation, which may play crucial roles in the carbon source transition from glucose to proteins, providing new potential therapeutic targets for metabolic disorders.

## Introduction

Under nutrient-deficient conditions, the proteome is remodeled through protein degradation and epigenetic alterations to adapt to environmental changes [Ref]^1,2^. A growing body of research indicates that amino acid starvation triggers the degradation of high-abundance proteins, such as Golgi proteins and ribosomal proteins, to supply energy [Ref]^3,4^. Previous studies have shown that glucose starvation (GS) drives increased glutamine entry into metabolic cycles [Ref]^5^. However, it remains unclear which proteins are degraded into glutamine to fuel metabolic pathways during glucose starvation.

H2Aub is one of the most abundant monoubiquitinated proteins in mammalian cells [Ref]^6^. The RING1B and BMI1 subunits of the PRC1 complex form a heterodimer that catalyzes mono-ubiquitination of vertebrate H2A at lysine 119 (K119) [Ref] ^7,8^. Recent studies reveal that PRC1 can also form liquid-liquid phase separation (LLPS) in the nucleus to mediate H2Aub [Ref]^9^. Glucose starvation reprograms transcriptional profiles of tumor-related genes by reducing the epigenetic marks H2Aub and H3ub [Ref]^10,11^. Our prior work suggests that Midnolin may provide amino acids to cells by mediating H2Aub degradation. While histone mono-ubiquitination is known to regulate gene expression epigenetically, whether it can be directly degraded in the nucleolus as an energy substrate remains an open question for further investigation.

Under energy stress, the degradation of proteins to recycle amino acids constitutes a fundamental biological process in mammalian cells [Ref]^12^. Major pathways for protein degradation include the ubiquitin-proteasome system and the autophagy-lysosome pathway [Ref]^13,14^. Lysosomes utilize acid hydrolases to break down macromolecules, such as proteins, nucleic acids, polysaccharides, and lipids, both inside and outside cells, thereby maintaining normal cellular metabolism and functional homeostasis [Ref]^15,16^. Autophagy includes macroautophagy, microautophagy, and chaperone-mediated autophagy [Ref]^17-19^. Nucleophagy, a selective form of autophagy, involves the encapsulation and degradation of nuclear components by autophagosomes, typically activated under stress conditions such as nutrient deprivation, oxidative stress, or DNA damage [Ref]^20-22^. For example, in yeast, nucleophagy proceeds through the formation of intranuclear membrane-derived vesicles, which deliver encapsulated nuclear material to lysosomes for degradation [Ref]^23,24^. In mammals, Atg39 acts as a critical nucleophagy receptor by interacting with Atg8 to promote the formation of nuclear-derived vesicles, facilitating nuclear membrane degradation [Ref]^25^. Nucleophagy plays vital roles in maintaining cellular homeostasis, stress adaptation, and longevity [Ref]^20,26,27^. However, whether lysosomes exist within the nucleus and how they function under starvation or other stress conditions remain unexplored.

In this study, we revealed that glucose starvation triggers H2Aub, a histone epigenetic modification, as a direct energy substrate within lysosome-like structures in the nucleolus. We further uncovered lysosome-like structures that degrade monoubiquitinated histones (e.g., H2Aub), closely linked to glucose and amino acid metabolism. Our previous AI-assisted screening revealed that Midnolin as a potential receptor mediating H2Aub degradation in nucleolar lysosomes. This work elucidates the molecular mechanism underlying the fundamental physiological process of carbon source switching from glucose to amino acids derived from protein degradation during glucose starvation, which may provide novel therapeutic avenues for metabolic disorders.

## Results

### Glucose starvation specifically mediates H2Aub downregulation

Previous studies have shown that glucose starvation induces amino acids to enter metabolic cycles for energy production, but the specific proteins degraded to generate these amino acids remain unknown. To identify proteins degraded for energy supply, we employed proteomics to quantify protein degradation during glucose starvation. In total. 7912 proteins were quantified (Supplementary Table1). We categorized protein abundance dynamics into four clusters (Fig. 1a). Cluster 1 proteins exhibited a rapid decrease immediately after glucose starvation treatment and were enriched in pathways related to chromatin assembly (Fig. 1b). Cluster 2 proteins showed delayed reduction only after 1 hour of glucose deprivation, suggesting their association with secondary responses to glucose starvation (Fig. 1a). These proteins were enriched in pathways such as cell cycle regulation, implying potential cell cycle arrest under glucose deficiency (Fig. 1b). Cluster 3 proteins displayed an immediate increase post-glucose starvation, indicating their role in regulating metabolic homeostasis under stress. These were significantly enriched in DNA damage response pathways (Fig. 1a). Cluster 4 proteins exhibited marked changes only after 1 hour, further supporting their involvement in secondary regulatory responses to glucose starvation (Fig. 1b).

**Figure 1:**
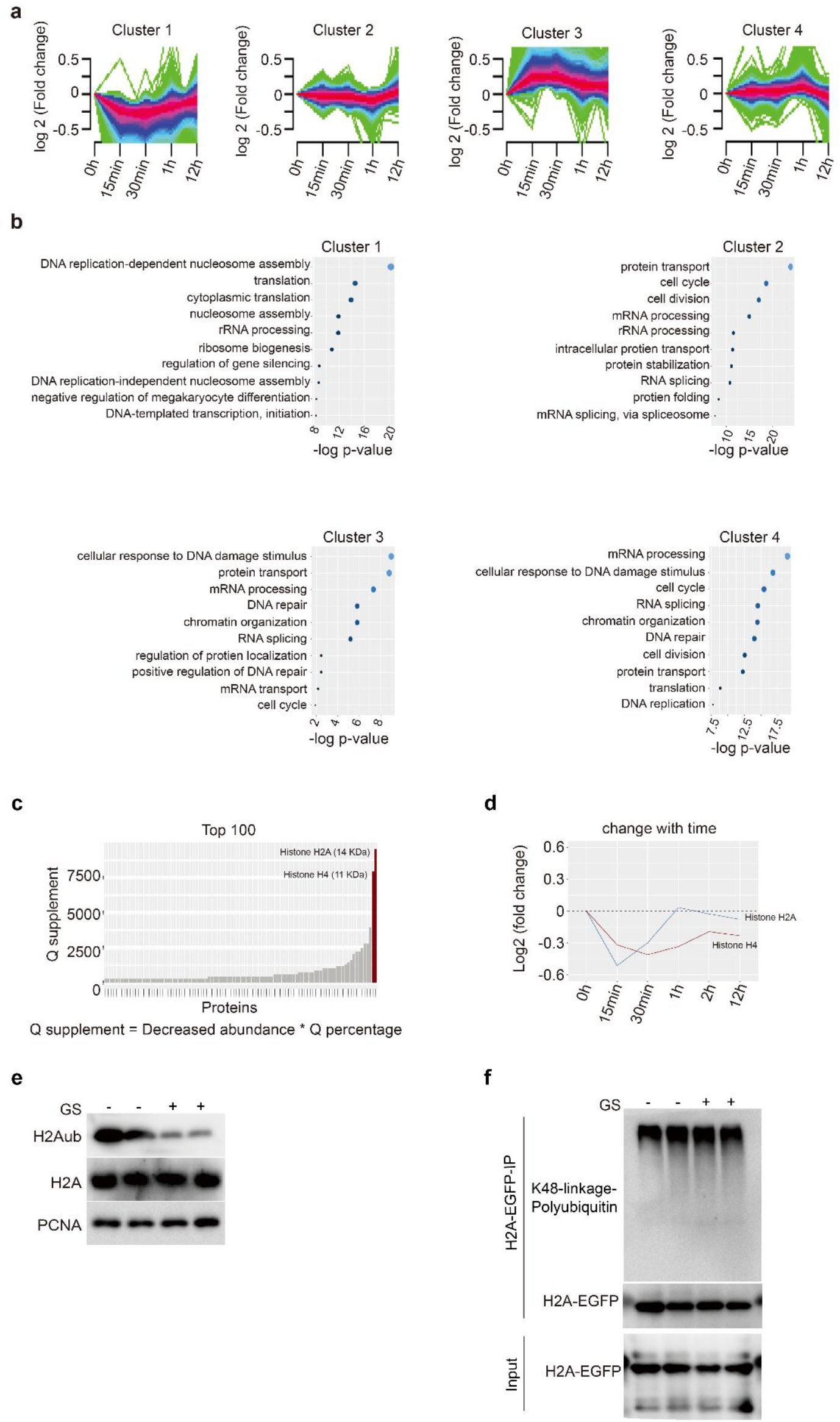
Under glucose starvation, monoubiquitinated histones (e.g., H2Aub) act as direct donors for degradation, with histones exhibiting the highest glutamine content among degradation products. a. Glucose starvation induces altered dynamics of protein degradation. b. Glucose starvation induces pathway enrichment of proteins with distinct dynamic patterns. c. Proteins are ranked by glutamine production capacity derived from reduced protein content. d. Mass spectrometry analysis reveals rapid depletion of histones H2A and H4 following glucose starvation. e. Glucose starvation does not affect total H2A levels but induces reduction of monoubiquitinated histone H2Aub content. f. Glucose starvation does not alter K48-linked polyubiquitination of total H2A.

To identify proteins that degrade to provide energy in the form of amino acids, we converted the reduced protein levels into estimated amino acid production. First, we calculated the fold reduction in protein abundance after 30 minutes of glucose starvation as -log_2_FC. Using the baseline proteomics abundance under untreated conditions as the reference, we then determined the top glutamine-supplying proteins after glucose starvation using the formula: Q supplement abundance = (abundance × (-log_2_FC)) × Q percentage, where Q percentage represents the proportion of glutamine residues within each protein’s total amino acid composition (Supplementary Table1). We observed that histones yield high glutamine levels, notably Histone H4 and H2A (Fig. 1c). Furthermore, Histone H4 and H2A exhibited a marked decline in abundance during early 30min glucose starvation (Fig. 1d).

To further validate histone abundance changes under glucose starvation, we performed Western blot analysis. We observed that glucose starvation did not significantly affect total histone; however, monoubiquitinated histones exhibited a marked reduction (Fig. 1e). Notably, glucose starvation did not induce an increase in K48-linked polyubiquitination of histones (Fig. 1f). These results indicate that glucose starvation selectively reduces the monoubiquitinated subset of histones, suggesting that monoubiquitinated histones, such as H2Aub, may serve as direct energy substrates through degradation.

### H2Aub was degraded via the lysosomal pathway

To investigate whether H2Aub is degraded via the proteasomal pathway, we treated cells with MG132, a proteasome inhibitor known to block proteasome activity and deplete intracellular amino acid pools. Surprisingly, MG132 not only failed to inhibit glucose starvation-induced H2Aub reduction but exacerbated H2Aub degradation more severely than glucose starvation alone, strongly suggesting that H2Aub may act as a direct amino acid donor for energy production (Fig. 2a). In contrast, the lysosomal pathway inhibitor chloroquine (CQ) effectively suppressed H2Aub reduction under glucose starvation (Fig. 2b). These findings further support that H2Aub is degraded through the lysosomal pathway to supply amino acids for energy metabolism.

**Figure 2:**
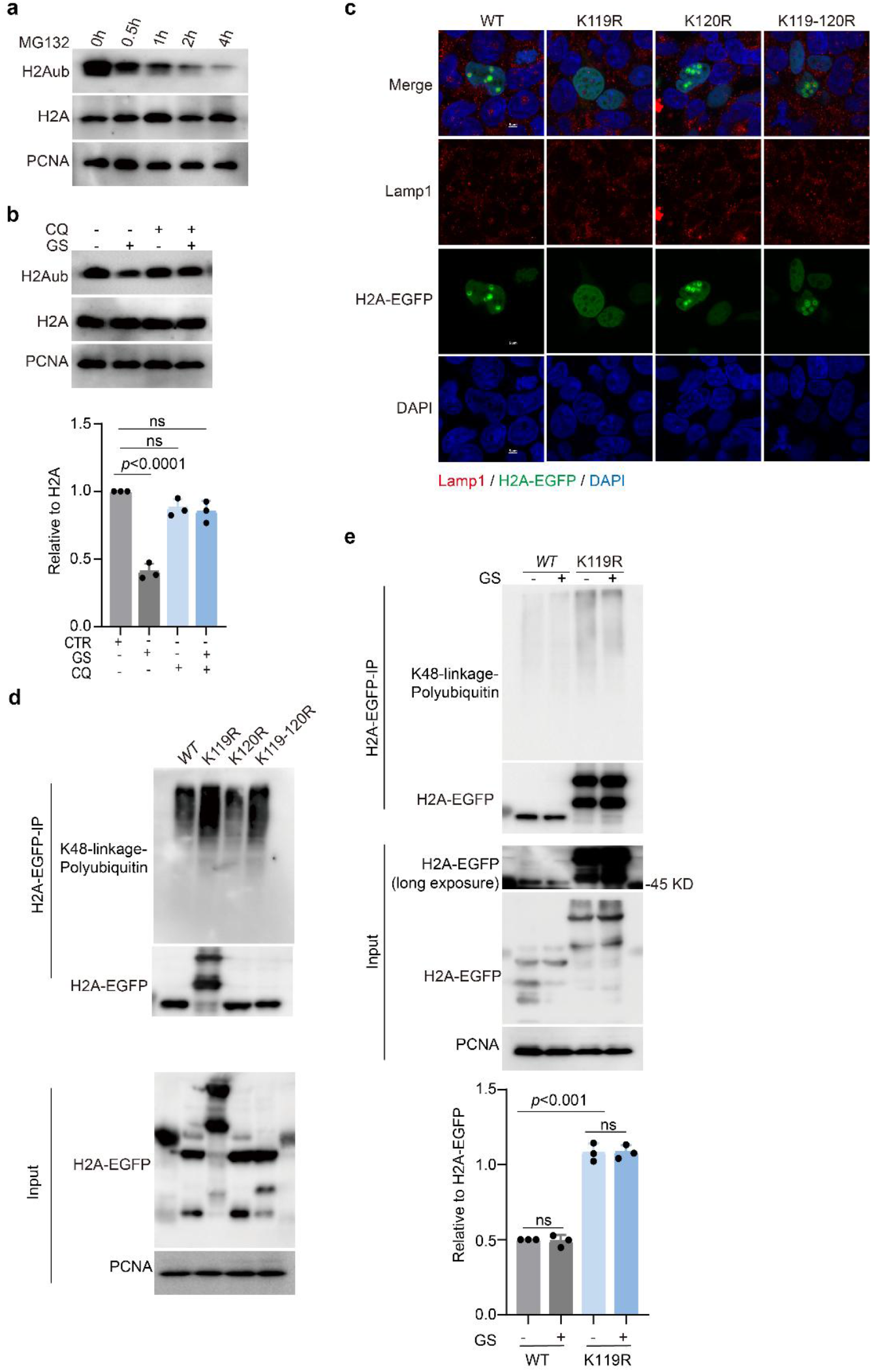
H2Aub undergoes degradation through the lysosomal pathway. a. MG132 treatment triggers rapid depletion of monoubiquitinated histone H2Aub. b. Western blot analysis and quantification show that CQ inhibits glucose starvation-induced reduction of H2Aub content. c. The co-localization of various H2A mutants with the lysosomal marker LAMP1 in the nucleolus. d. Distribution of monoubiquitination and K48-linked ubiquitination in various H2A-EGFP mutants. e. Western blot analysis and quantification show that the monoubiquitination site K119 of H2A is essential for glucose starvation-induced degradation of H2A ubiquitination (H2Aub).

Consistent with the lysosomal degradation of H2Aub, overexpression of the histone H2A plasmid revealed concentric colocalization with lysosome marker LAMP1 in the nucleolus (Fig. 2c). However, when the mono-ubiquitination site K119 was mutated to arginine (K119R), H2A failed to localize to the nucleolus due to the loss of mono-ubiquitination (Fig. 2c). Intriguingly, the K119R mutation triggered extensive polyubiquitination at the adjacent K120 residue. This aberrant polyubiquitination was abolished in the double mutant (K119R/K120R), which lacks robust polyubiquitination signals. Wild-type (WT), K120R, and K119R/K120R mutants—all devoid of the excessive polyubiquitination caused by K119R—retained nucleolar accumulation (Fig. 2c), demonstrating that H2A polyubiquitination is excluded from the nucleolar targeting. Furthermore, under glucose starvation, the K119R mutation abolished changes in ubiquitinated H2A (H2Aub) levels (Fig. 2d). These findings conclusively establish that nucleolar degradation of H2Aub depends on its mono-ubiquitination at the K119 site.

### lysosome-like structures in the nucleolus regulate the balance of carbon source

To further elucidate the role of nuclear lysosome-like structures (LLS) in the carbon source transition from glucose to amino acids, we examined Lamp1 dynamics under glucose starvation. Glucose starvation induced significant foci formation of Lamp1 (Fig. 3a). Similarly, MG132 promoted the clustering of lysosome-like structures within the nucleus (Fig. 3a), whereas chloroquine (CQ) alone had no observable effect. However, combined treatment with glucose starvation and CQ resulted in the formation of massive Lamp1 foci in the nucleolus (Fig. 3a), further supporting H2Aub degradation via the nucleolar lysosomal pathway. Additionally, prolonged culture without media replacement led to progressive LLS aggregation, transitioning from dispersed nucleolar localization to 1-2 prominent foci (Fig. 3b). These observations demonstrate that nuclear LLS are highly sensitive to nutrient availability.

**Figure 3:**
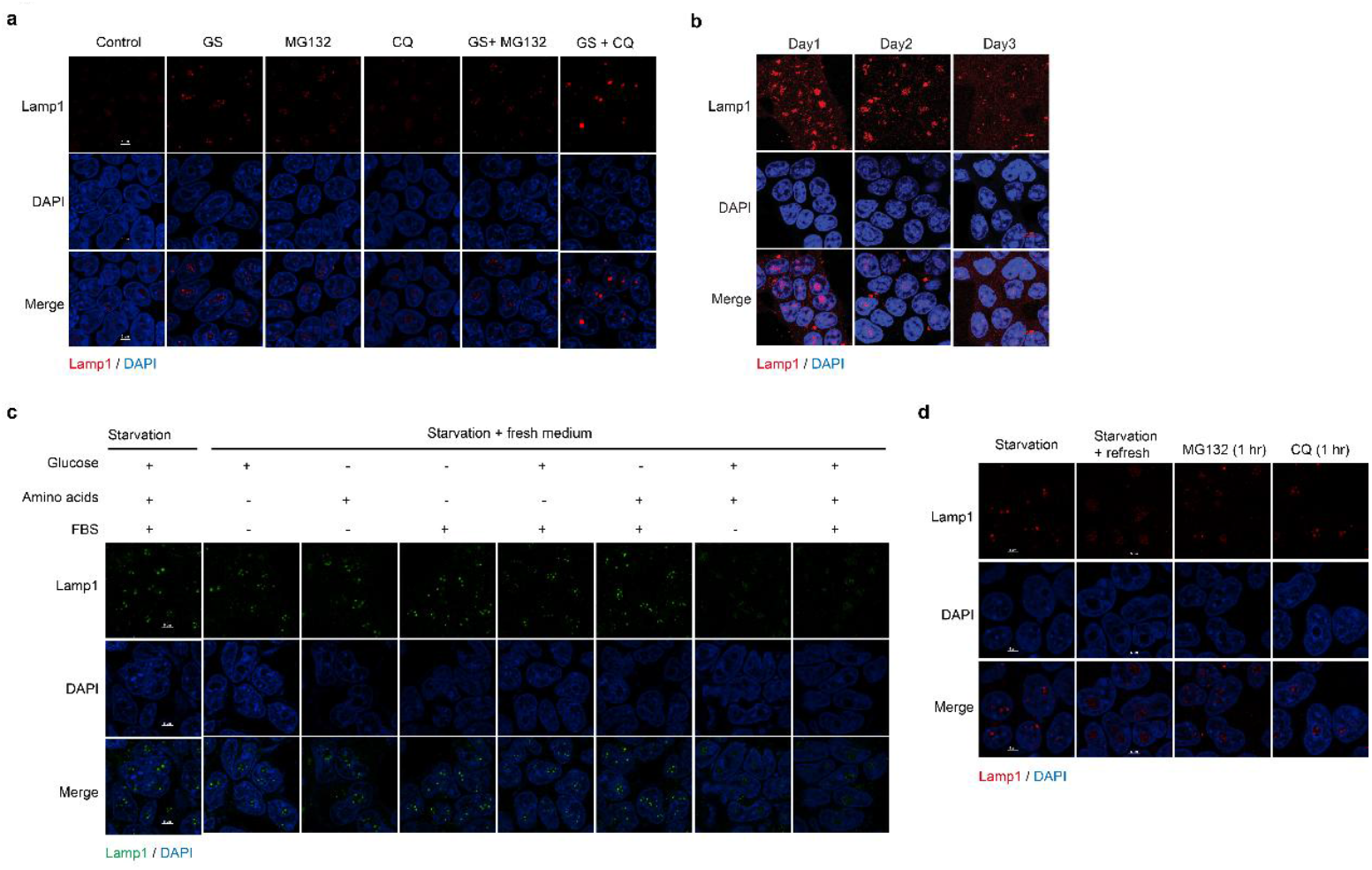
Lysosome-like structures within the nucleolus mediate the metabolic shift of carbon source utilization from glucose to protein catabolism. a. Lysosome-like marker Lamp1 forms massive foci in the nucleolus exclusively under combined glucose starvation and CQ treatment. Endogenous Lamp1 (red) were stained with rabbit anti-Lamp1 antibody, and were imaged by confocal microscopy. b. Prolonged starvation also induces large Lamp1 foci within the nucleolus. c. Rapid dissolution of Lamp1 foci occurs only in the simultaneous presence of glucose and amino acids. d. MG132 exhibits no significant effect on Lamp1 aggregate dissolution, whereas CQ delays the disaggregation process.

To further demonstrate that nucleolar Lamp1 mediates the carbon source switching between glucose and amino acids, we performed nutrient replenishment experiments after 48 hours of starvation. Individual nutrients—such as fetal bovine serum (FBS), amino acid mixture, or glucose alone—failed to disperse Lamp1 foci. Similarly, combinations of two nutrients (e.g., glucose + FBS or amino acids + FBS) also did not induce Lamp1 disassembly. Only when both glucose and amino acids were provided simultaneously did Lamp1 foci undergo complete dispersion (Fig. 3c). These results conclusively establish Lamp1’s role in mediating carbon source switching between glucose and amino acids.

To investigate Lamp1 disassembly, fresh medium was supplemented with MG132 or CQ. We observed that MG132 had no significant impact on Lamp1 disassembly, whereas CQ partially inhibited this process (Fig. 3d). These findings further confirm that Lamp1 functions as a lysosome-like structure within the nucleus, actively participating in energy metabolism.

### Midnolin acts as a receptor that mediates H2Aub degradation to regulate the cell cycle

While our findings suggest that nucleolar lysosomal pathways may degrade monoubiquitinated histones to supply energy, the receptor mediating H2Aub degradation remained unknown. We previously identified Midnolin as a candidate receptor through AI-assisted screening. Prior studies report Midnolin’s role in degrading ubiquitinated proteins; here, we demonstrate that Midnolin also facilitates lysosomal degradation of monoubiquitinated histones. Notably, nucleolar Midnolin aggregation increased upon GS treatment, and Midnolin exhibited strong interaction with lysosome-like structures (LLS) in the nucleolus (Fig. 4a).

**Figure 4:**
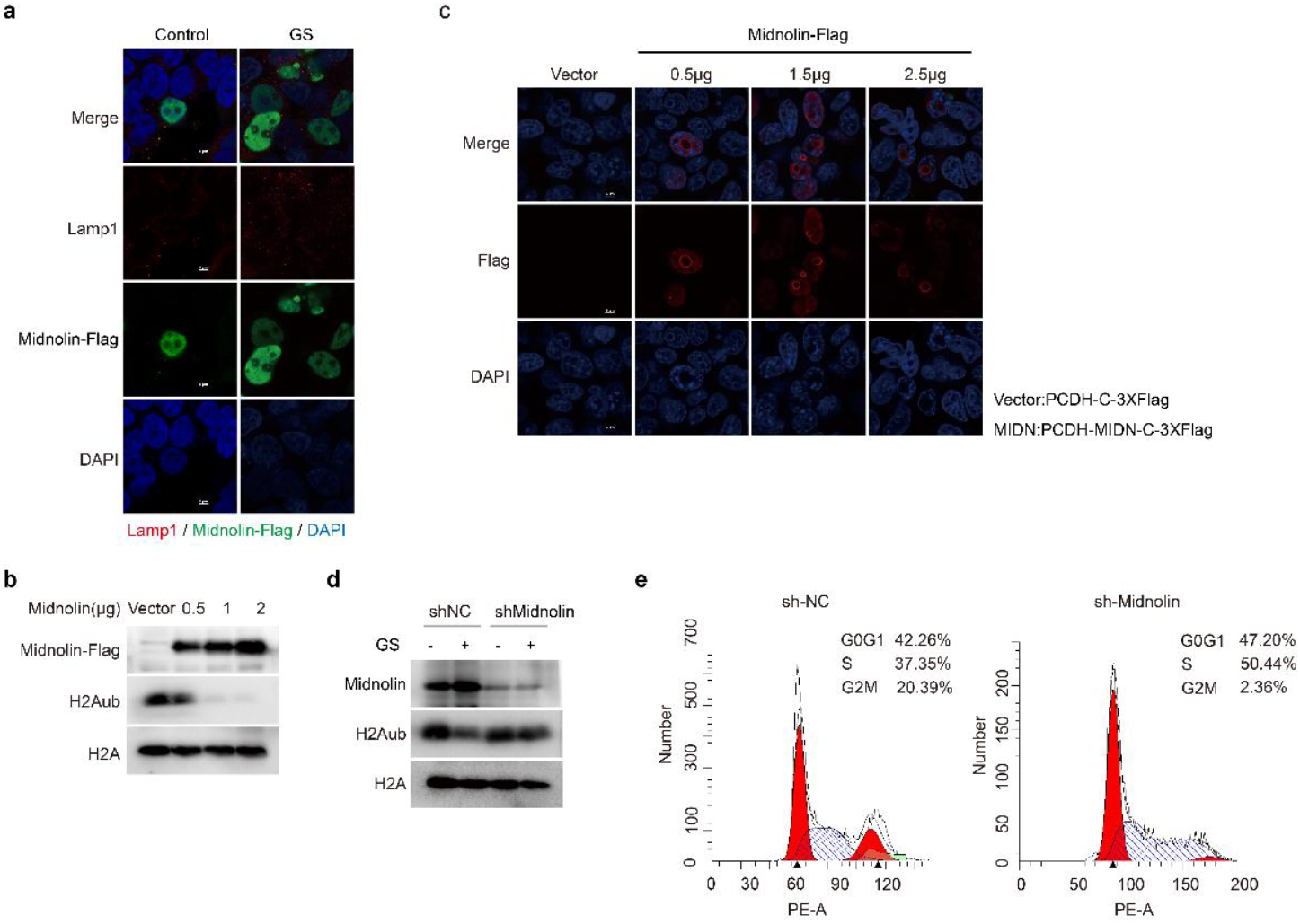
Midnolin in the nucleolus acts as a receptor mediating the degradation of H2Aub within nucleolar lysosome-like structures. a. Glucose starvation induces increased aggregation of Midnolin in the nucleolus. Midnolin-Flag were transfected into 293T. Then cells were treated with glucose starvation. Endogenous Lamp1(red) and overexpressed Midnolin-Flag (green) were stained with rabbit anti-Lamp1 antibody and mouse anti-Flag antibody respectively, and were imaged by confocal microscopy. b. Midnolin overexpression reduces H2Aub levels. c. Midnolin overexpression localizes to the nucleolus. Overexpressed Midnolin-Flag (red) were stained with mouse anti-Flag antibody, and were imaged by confocal microscopy. d. shMidnolin inhibits glucose starvation-induced degradation of H2Aub.

Furthermore, overexpression of Midnolin promoted H2Aub degradation (Fig. 4b), accompanied by the formation of distinct annular structures within nucleoli (Fig. 4c). Conversely, Midnolin knockdown effectively blocked the GS-induced reduction of H2Aub levels (Fig. 4d). These results further confirm that Midnolin mediates H2Aub degradation within the nucleolus.

Proteomic quantification revealed significant alterations in cell cycle-related proteins under glucose starvation, prompting us to hypothesize that energy-supplying protein degradation during glucose deprivation might regulate cell cycle progression. Indeed, Midnolin depletion led to S-phase arrest, demonstrating that this proteolytic energy-generating process critically controls cell cycle dynamics (Fig. 4e).

## Discussion

Previous studies indicate that mammals degrade proteins to supply energy under nutrient stress and adapt to environmental changes through epigenetic regulation of gene expression [Ref]^28,29^. Here, we demonstrate that monoubiquitinated histone H2Aub, traditionally regarded as an epigenetic modifier, serves as a direct energy substrate, undergoing degradation in the nucleolus to release amino acids for energy production. Quantitative proteomics revealed that glucose starvation triggers histone degradation, with glutamine emerging as the most abundantly generated amino acid. We further identified lysosome-like structures (LLS) within the nucleolus, likely mediating H2Aub degradation to fuel energy metabolism. Through functional screening, we established Midnolin as a novel receptor responsible for targeting H2Aub to nucleolar LLS for degradation.

Previous studies have shown that nutrient deprivation enables adaptation to environmental changes through epigenetic reprogramming of gene expression. For instance, glucose starvation induces acetylome change, facilitating the expression of gluconeogenic genes [Ref]^30,31^. Here, we demonstrate that histone H2Aub, one the most abundant monoubiquitinated protein in cells, not only participates in gene expression regulation [Ref]^32-34^ but also acts as a direct energy substrate, undergoing degradation in the nucleolus to supply amino acids for energy production. Quantitative proteomics revealed that histone degradation under glucose starvation predominantly generates glutamine. Furthermore, lysosome-like structures (LLS) within the nucleolus likely mediate H2Aub degradation to fuel energy metabolism. Through screening, we identified Midnolin as the receptor responsible for targeting H2Aub to nucleolar LLS for degradation. These findings suggest that histones, as highly abundant proteins, dually function as epigenetic carriers and energy reservoirs, aligning with the rapid responsiveness of epigenetic mechanisms to environmental stress. Other monoubiquitinated histones, such as H2Bub, H3ub, H4ub, may undergo similar adaptive changes [Ref]^35,36^, though this requires further investigation.

While the existence of lysosome-like structures (LLS) within the nucleolus has not been previously reported, earlier studies have observed similar structures without characterizing their functional relevance [Ref]^37^. In our study, we unexpectedly identified LLS in the nucleolus and demonstrated their involvement in degrading monoubiquitinated histones. Through systematic treatments, we found that LLS signals dynamically respond to nutrient availability and pharmacological perturbations, exhibiting heightened sensitivity to glucose and amino acid metabolic states. These observations strongly suggest that nucleolar LLS participate in the metabolic shift from glucose utilization to amino acid generation via protein degradation. However, since LLS predominantly localize to the cytoplasm, overexpression of LLS-associated proteins only allowed detection of cytoplasmic LLS by immunofluorescence, likely due to the low abundance of nuclear LLS. To visualize nucleolar LLS, future studies should investigate potential structural or functional distinctions between nucleolar and cytoplasmic LLS that underlie their differential subcellular distribution.

Midnolin has been reported as a nuclear protein that recognizes ubiquitinated substrates and mediates their proteasomal degradation [Ref]^38,39^. However, through data mining and validation, we identified a long isoform of Midnolin localized to the nucleolus, where it exhibits strong colocalization with nucleolar proteins. Notably, glucose starvation enhanced its nucleolar colocalization. Whether and how the previously studied short Midnolin isoform contributes to monoubiquitinated histone H2Aub degradation remains an open question requiring further investigation [Ref]^40,41^.

Tumor cells exhibit exceptionally high energy demands, with elevated levels of the mono-ubiquitinating enzyme BMI1, responsible for H2A mono-ubiquitination, being a recognized hallmark of malignancies [Ref]^42-45^. Notably, we found that knockdown of Midnolin, a key protein in the H2Aub degradation pathway, significantly arrests the cell cycle of tumor cells. These findings suggest that modulating H2Aub levels may restrict the energy available for tumor cell proliferation, effectively starving tumor cells and thereby suppressing tumor growth.

## Methods

### Cell Culture and Transfections

Cells were maintained in high-glucose DMEM medium (Saivei, G4511-500ML) supplemented with 10% fetal bovine serum (FBS) (Novizan, F101) and 1% penicillin-streptomycin (Zhongqiao Newchuan, CSP006) in a 37°C incubator with 5% CO_2_. Twenty-four hours before transfection, cells were seeded into 6-well plates using complete growth medium. For Lipofectamine 2000 (Biosharp, BL623B)-mediated transfection, DNA-lipid complexes were prepared according to the manufacturer’s protocol. The complexes were added to cells cultured in serum-free medium, and the plates were gently rocked to ensure uniform distribution. Following a 6-hour incubation, the medium was replaced with fresh complete growth medium. Cells were transfected for 24–48 hours, after which they were treated with glucose-free medium or 20 μM MG132 as required by the experiment.

### The product numbers (Catalog numbers) of the antibodies used are as follows

Ubiquityl-HistoneH2A(Lys119) (D27C4) (CellSignalingTechnology, 8240S); K48-linkagePolyubiquitin (D9D5) Rabbit mAb (CellSignalingTechnology, 8081S); Midnolin Rabbit Polyclonal Antibody(abbiotec, 251274); GFP tag Polyclonal antibody (Proteintech, 50430-2-AP); PCNA Rabbit pAb (ABclonal, A1336); HRP-Labeled Goat Anti-Rabbit IgG (ZSGB-BIO, ZB-2301); Anti-Flag Magnetic Beads (Beyotime, P2115-2ml); Anti-GFP Magnetic Beads (Beyotime, P2132-2ml); Mouse IgG Magnetic Beads (Beyotime, P2171-1ml); FLAG mouse (Merck, F1804); LAMP1 Lysosome Marker (Abcam, ab24170); Alexa Fluor 594 Goat anti-Rabbit IgG (ThermoFisher, A-110037); Alexa Fluor 488 Goat anti-Mouse IgG(ThermoFisher, A-11029); DAPI solution (Solarbio, C0060).

### Immunofluorescence (IF)

At room temperature, cells were fixed with 4% paraformaldehyde (Beyotime, P0099-100ml) for 15 minutes, followed by permeabilization with 0.2% Triton X-100 (Selleck, T8200) for 10 minutes. Cells were then blocked with 1% BSA (Beyotime, ST2249) at room temperature for 30 minutes. After blocking, the cells were incubated with the primary antibody overnight at 4°C. Following washing, the cells were incubated in the dark at room temperature for 1–2 hours with a mixture of secondary antibodies (Alexa Fluor 488 or Alexa Fluor 594, 1:200) and DAPI (1 mg/mL, 1:200 working concentration). After washing, the cells were mounted and imaged using a Nikon confocal laser scanning microscope.

### Immunoprecipitation (IP)

Cells were transfected into 100 mm culture dishes and supplemented with 800 μL IP Lysis Buffer (Biosharp, BL509A). Sonication was performed 15 times (each for 2-3 seconds). Subsequently, the mixture was centrifuged at 4°C and 12,000 rpm for 10 minutes. The supernatant containing the lysate was incubated with 10 μL mouse IgG magnetic beads (Beyotime, 2171-1ml) on a rotating wheel at 4°C for 15 minutes to perform pre-clearing. After magnetic separation, 20 μL Anti-Flag magnetic beads (Biosharp, P2115-2ml) were added to the supernatant, and the mixture was incubated on a rotating wheel at 4°C overnight. The beads were washed three times with IP Lysis Buffer (5 minutes each wash). The beads were resuspended in 50 μL of 1× SDS loading buffer after removing the wash solution.

### Western blotting (WB)

Cells were digested and collected, and then lysed using a lysis buffer containing the following components: 50 mM Tris/HCl (pH 7.5) (Biosharp, ST768), 0.5% Nonidet P-40 (Selleck, N8030), 1 mM EDTA (Biosharp, ST066), 150 mM NaCl (Aladdin, G8200), 1 mM dithiothreitol (DTT) (Biosharp, ST403-5g), 1 mM phenylmethylsulfonyl fluoride (PMSF) (Biosharp, C0351-1ml), and 10 mM Complete protease inhibitor cocktail (Targetmol, M1500). Clarified cell lysates with a total protein concentration of 30-50 μg were collected for subsequent Western blot analysis.

### Knockdown Experiments

Midnolin,shRNAs were purchased from Kingbrave Biotech (Wuhan). The lentiviraltarget sequences were shMidnolin #1: 5’-GATGTGAACATCACGTGTTAT-3’, shMidnolin #2: 5’-TCATTATGCACTTTACAGAT-3’. Cells were selected with 2 μg/ml puromycin (Biosharp, C0351-1ml) for one week to obtain lentivirus-positive cells. Subsequently, the selected cells were validated by Western blot analysis.

### Cell Cycle Analysis by Flow Cytometry

Trypsin (VivaCell, C3530-0500) was used to digest and collect adherent cells, which were then washed with PBS. Cells were fixed with cold ethanol pre-cooled to -20°C at 4°C for at least 4 hours or overnight. The cells were stained with PI/RNase staining solution (RNase to eliminate RNA interference), incubated at 37°C in the dark for 30 minutes. Flow cytometry was performed to analyze the samples, and professional software was used to fit the cell cycle distribution data.

### Site-Directed Mutagenesis

The template plasmid with GC content between 40-55% was selected to achieve optimal mutation efficiency. Site-specific mutagenesis was performed according to the manufacturer’s protocol of the site-directed mutagenesis kit (Biosharp, D0206M). The reaction products were directly transformed into DH5α competent cells, and plated onto LB agar plates containing the respective antibiotics. Single colonies were picked for mini-plasmid preparation, and the success of the mutation was verified by sequencing or restriction digest.

### Sample preparation for LC-MS analysis

The 293T cells were subjected to glucose starvation treatment for 0 hours, 15 minutes, 30 minutes, 1 hour, and 12 hours respectively. Subsequently, cells were harvested using urea lysis buffer containing 8 M urea, 100 mM Tris-HCl (pH 8.5), 150 mM NaCl, and protease inhibitor cocktail (Targetmol, M1500). The lysates were centrifuged at 10,000 rpm for 15 minutes, and the protein concentration in the supernatant was determined using the BCA protein assay (Pierce, USA). Proteins were digested with trypsin (promega, V5113) at a 1:100 ratio. The samples from the control group are used for measuring protein reference abundance. Then, wo biological replicate samples were set up for each time point and labeled using TMT reagents (Thermo Fisher Scientific) according to the manufacturer’s instructions. The brief procedure was as follows: 100 ug of digested peptides were resuspended in 100 ul of 100 mM TEAB buffer and labeled with distinct TMT reagents. Finally, all TMT-labeled peptides were equally pooled for subsequent analysis.

### Basic reversed phase (RP) chromatography

For TMT labeling sample analysis, the offline basic reversed-phase high-performance liquid chromatography (bRP-HPLC) fractionation was performed using an Agilent 1260 HPLC system equipped with an XBridge BEH C18 column (1 mm,150 mm,130 Å, 3.5 μm, Waters, Herts, UK). Peptides were fractionated through a 72-min bRP-HPLC gradient under constant flow rate of 70 μl/min throughout the mobile phase elution. A total of 49 fractions were collected into 1.5 mL microcentrifuge tubes (Axygen, N/A)) after injection. These minor fractions were subsequently non-consecutively pooled into 7 major fractions and completely dried using a vacuum concentrator. The resulting peptides were finally reconstituted in aqueous solution containing 0.1% formic acid (FA) for subsequent liquid chromatography-mass spectrometry (LC-MS) analysis.

### LC–MS Data Acquisition

The online liquid chromatography-tandem mass spectrometry (LC-MS/MS) analysis was performed on a Tribrid Orbitrap Fusion mass spectrometer coupled with a NanoLC-1000 high-performance liquid chromatography system (Thermo Fisher Scientific). Peptide separation was achieved using a self-packed 15 cm fritless column (C18 packing material, 1.9 μm particle size; Dr. Maisch GmbH, Germany) at a flow rate of 300 nl/min. For TMT-labeled samples, the mass spectrometer operated in data-dependent acquisition mode: full MS1 scans were first acquired in the Orbitrap (m/z range 350-1500; resolution 60,000; automatic gain control [AGC] target 400,000), followed by MS2 scans (normalized collision energy of 37% with 5% stepped collision energy; resolution 30,000; AGC target 50,000; maximum injection time 60 ms). The isolation windows were set to 1.5 m/z and 1.6 m/z for TMT and SILAC-labeled precursors, respectively. A dynamic exclusion duration of 45 s was applied for precursor ion selection.

### Data Search and Post analysis

The raw data from different fractions were jointly searched against the UniProt human database using Proteome Discoverer 2.1 (PD 2.1) software. The precursor mass tolerance and fragment ion mass tolerance were set to 10 ppm and 0.1 Da, respectively. Enzymatic digestion parameters were specified as trypsin with a maximum of three allowed missed cleavages. For protein reference analysis, carbamidomethylation and methionine oxidation was set as static and variable modifications respectively. For TMT sample analysis, static modifications included carbamidomethylation of cysteine (+57.02 Da) and 10-plex TMT labeling (+229.16 Da) on lysine residues and peptide N-termini, while dynamic modifications permitted methionine oxidation (+15.99 Da). A false discovery rate (FDR) threshold of 0.01 was applied at the peptide level, defined as the ratio of false positive identifications to total identifications. The intensities of all reporter ion channels were initially normalized using median values. Protein expression fold changes were derived from the ratios of average reporter ion intensities, and corresponding p-values were calculated using Student’s t-test to assess the statistical significance of differentially expressed proteins.

## Acknowledgements & Fundings

This research was funded by grants from National Natural Science Foundation of China (32400978) and Start-up Grant for Yellow River Scholar at Henan University. We thank Chensong Zhang for helpful discussion and advice on this project.

## Authors contribution

LY, and YQL conceived the project and designed the study. LY, JLJ, SYM, JS, HHL, YXL, YY and NF conducted experiments. LY, YQL, JLJ, SYM, JS, HHL, YXL interpreted data. LY and YQL contributed to the discussion and critical regents. LY and YQL wrote the paper.

